# Nanoscale infrared spectroscopy identifies parallel to antiparallel beta sheet transformation of Aβ fibrils

**DOI:** 10.1101/2022.09.27.509729

**Authors:** Siddhartha Banerjee, Divya Baghel, Md. Hasan-ul-Iqbal, Ayanjeet Ghosh

## Abstract

Spontaneous aggregation of amyloid beta (Aβ) proteins leading to the formation of oligomers and eventually into fibrils has been identified as a key pathological signature of Alzheimer’s disease. Structure of late stage aggregates have been studied in depth by conventional structural biology techniques including Nuclear Magnetic Resonance, X-ray crystallography and Infrared Spectroscopy; however the structure of early-stage aggregates is less known due to their transient nature. As a result, the structural evolution of amyloid aggregates from its early oligomers to mature fibril is still not fully understood. Here we have applied AFM-IR nanospectroscopy to investigate the aggregation of Aβ 16-22, which spans the amyloidogenic core of the amyloid beta peptide. Our results demonstrate that Aβ 16-22 involves a structural transition from oligomers with parallel beta sheets to antiparallel fibrils through disordered and possibly helical intermediate fibril structures, contrary to the known aggregation pathway of full-length Aβ.

## Introduction

The most common pathological hallmark of the Alzheimer’s disease is the presence of plaque found in the brain^1^. These extracellular plaques are mainly composed of aggregates of different isomers of the amyloid beta (Aβ) peptide^1–3^. Structure of amyloid protein aggregates, specifically fibrils, which are generated at the late stage of aggregation, have been widely investigated, and amyloid fibrils are known to have in-register parallel beta sheet organization^4–7^, although N-terminal of the peptide remains disordered even in the fibrillar stage^5, 6, 8^. The 17-21 residue region of the Aβ sequence constitutes a hydrophobic core that is known to be pivotal for aggregation of the full-length peptide^7^. Interestingly, small peptides containing this sequence have been shown to have an inhibitory effect on the aggregation of full-length Aβ^9^. As a result, several studies have aimed to understand the structural evolution and aggregation of this core peptide sequence, and it has been shown that short peptides spanning this sequence typically form highly ordered fibrils with antiparallel beta sheet structure^10, 11^. Similar to the larger isoforms of Aβ, while the structure of the fibrillar aggregates of the hydrophobic core containing peptides is known, the early-stage aggregates which transition to antiparallel fibrils are significantly less understood. Given that full-length Aβ forms fibrils with parallel beta sheet structure, the structural information of early stage Aβ aggregates cannot necessarily be applied to these shorter peptides. This lack of knowledge exists because common structural biology tools, namely Nuclear Magnetic Resonance (NMR) and infrared (IR) spectroscopies work well on conformationally homogeneous specimens but cannot distinguish between structures at a single aggregate level. Amyloid aggregation produces variety of aggregated species, some of which can be short lived and undergo spontaneous transformation into more stable structures, making it challenging to follow the structural reorganization.

In this report, we use atomic force microscopy (AFM) combined with infrared spectroscopy (AFM-IR) to address this issue. AFM provides nanoscale morphological information of the aggregates, whereas IR spectroscopy reveals the secondary structure of them at individual aggregate level^12, 13^. We have applied AFM-IR to investigate the structural reorganization of the Aβ 16-22 peptide, which is the smallest peptide spanning the hydrophobic core that forms fibrils^10^. The late-stage fibrils of Aβ 16-22 have been previously shown to have anti-parallel beta structure; we show that early-stage aggregates, including both prefibrillar aggregates and fibrils, adopt an ordered parallel beta structure which transforms into anti-parallel beta structure upon fibril maturation.

## Results and discussion

To understand the structural changes of Aβ 16-22, we initiated aggregation by incubating 1mM protein in 10 mM HCl at 37°C without any agitation and monitoring them after 15 min, 2 h, 6 h and 24 h of incubation. Fig. 1A shows the oligomers which are generated within 15 minutes of aggregation. Oligomers appeared to be mostly globular in shape with different sizes. IR spectra of the amide-I region, which is reflective of protein secondary structure^14^, were consequently recorded on individual oligomers. Locations from where the spectra were recorded are shown in Supporting Information (Fig. S1). The observed spectra can be essentially categorized into two subtypes: the first type shows a dominant peak at 1636 cm^−1^ (Fig. 1B) with smaller shoulders at ~1666 cm^−1^ and 1690 cm^−1^; the second type exhibits the same peaks, but the peak at 1690 cm^−1^ (Fig. 1D). The second derivatives of the average spectra of each subtype are shown in Figures 1C and E, which confirm the presence of three underlying bands for each. Representative spectra from different oligomers are shown in the Supporting Information (Fig. S2). This spectral heterogeneity clearly indicates that at the very early stage of aggregation, oligomers with two different secondary structures are formed. We discuss the structural implications of these spectra later.

**Figure 1:**
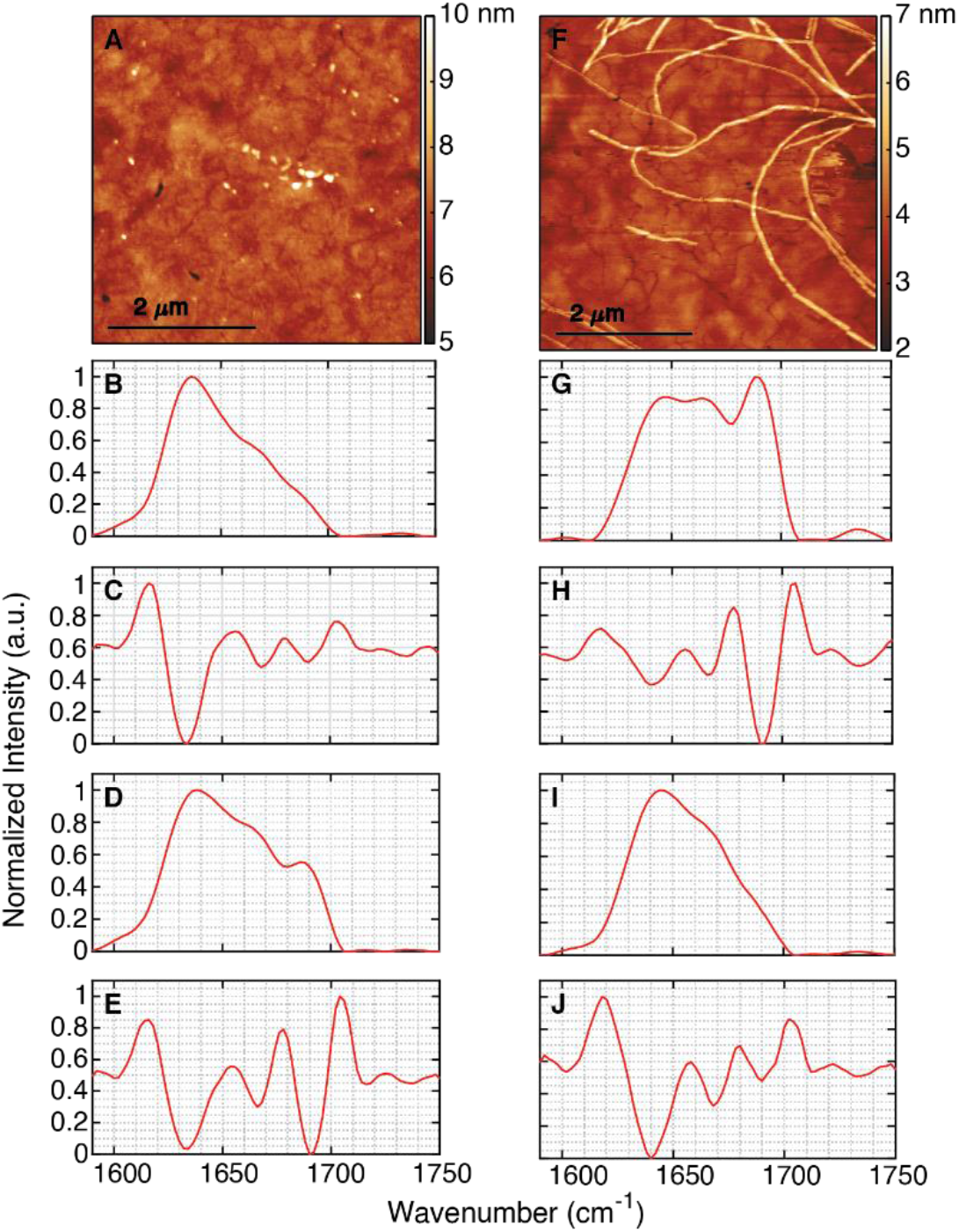
AFM-IR characterization of Aβ 16-22 oligomers and early-stage fibrils in 10 mM HCl. (A) AFM topography of oligomers after 15 min incubation. (B-C) Average IR spectra and second derivative of the first spectral subtype seen in oligomers. (D-E) Average IR spectra and second derivative of the second spectral subtype. (F) AFM image of fibrils generated after 2 h of incubation. (G-H) Average IR spectra and second derivative of the first spectral subtype seen in fibrils. and (I-J) Average IR spectra and second derivative of the second fibrillar subtype.

After 2 h of incubation fibrillar aggregates were observed (Fig. 1F). IR spectra were acquired along the same fibril and also on different fibrils (Fig. S1). Here also we observe mainly two types of amide-I bands; however, the spectral differences are not from different fibrils: the same fibril exhibits significant variations in spectra. One spectral type is very similar to the first observed in oligomers (Fig. 1B), however, the peak position exhibits a shift to 1644 cm-1 (Fig. 1I and Fig. S2). The second type of amide- I band is significantly different from oligomeric spectra, exhibiting a peak at 1690 cm-1 and two shoulders at 1644 cm-1 and 1676 cm-1 (Fig. 1G). The positions of the sub-bands are verified by the spectral second derivatives, shown in Figures 1H and J. Representative spectra from different spatial locations are shown in the Supporting Information (Fig. S2).

After 6h, we observed the transformation of individual fibrils into large fibrillar networks (Fig. 2A). The average IR spectrum, from different points along the fibrillar network, are shown in Fig. 2B and Fig. S3. Unlike the earlier fibrils, the spectra are homogeneous; the amide-I band shows two peaks at 1628 cm^−1^ and 1690 cm^−1^, where the relative peak intensity at 1628 cm^−1^ is higher compared to 1690 cm^−1.^ The spectra also exhibit weak bands between 1650-1660 cm^−1^; these can be clearly seen in the second derivative spectra (Fig. 2C). Fibril networks became denser after 24h of aggregation, and layers of fibrils were observed in the AFM topograph (Fig. 2D). As before, IR spectra were acquired at multiple spatial locations and the mean spectrum is shown in Figure 2E along with the second derivative (Fig. 2F). Similar to the 6h fibrils, the spectra do not exhibit heterogeneity, and only one type of spectrum is observed, containing two discrete peaks at 1628 cm^−1^ and 1690 cm^−1^.

**Figure 2:**
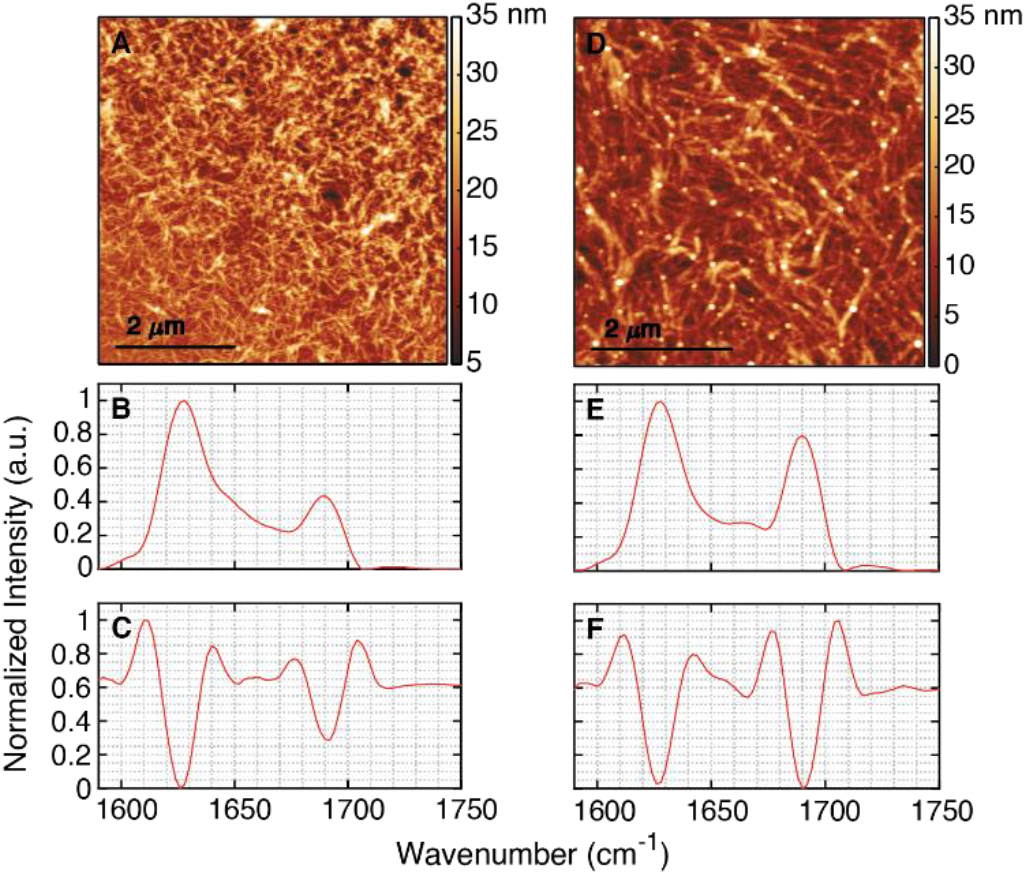
Structural investigation of Aβ 16-22 fibrils obtained in 10 mM HCl. (A) AFM image of fibrils obtained after 6h of incubation. (B) Average IR spectrum and (C) its second derivative from 6 h sample. (D) AFM image of fibrils generated after 24 h of incubation. (E) Average IR spectrum and (F) its second derivative from 24 h sample.

The aggregation pathway of the full length Aβ peptide has been studied extensively, and it is understood that the morphological transformation is accompanied by a change in secondary structure, specifically from antiparallel beta sheets in oligomers to parallel beta sheets in fibrils. However, the precise aggregation pathway leading to antiparallel fibrils for Aβ 16-22 is not known. Extrapolation of the aggregation mechanism of the full-length peptide would suggest maturation of antiparallel oligomers to antiparallel fibrils. Instead, we find two distinct structural states for oligomers (Fig. 1) and early-stage fibrils (Fig. 2), which are morphologically identical. It should be noted that this does not reflect polymorphism: we do not observe different morphologies of aggregates with different secondary structure; instead, we observe oligomers and early-stage fibrils that are different with respect to protein secondary structure, exhibiting no significant variations in morphology. The 2h fibrils in fact show spectral variations along the same fibril, which clearly indicates that we do not observe distinct polymorphs. We have recently reported similar observations in Aβ42^13^, and these results point to heterogeneity being a key structural facet of early-stage amyloid aggregates. Both types of oligomers exhibit the same spectral bands or substructures, but differ significantly in terms of their relative intensities and thus in terms of the overall secondary structure. Based on reported characteristic absorptions corresponding to different secondary structures^14^, the 1636 cm^−1^ band can be assigned to beta sheets. The high wavenumber band at 1690 cm^−1^ also corresponds to beta structure, specifically to antiparallel beta sheets, whereas parallel beta sheets only contain the low wavenumber band. This suggests that both polymorphs contain beta sheets: either a mixture of parallel and antiparallel, or antiparallel only. Both polymorphs also exhibit a band at 1666 cm^−1^, which likely arises from disordered/non beta sheet conformations. This peak can also correspond to a beta turn; but the disordered structure is a more plausible explanation for a small heptapeptide like Aβ 16-22. The relative intensity of the 1690 cm^−1^ peak is significantly higher than the other bands in one of the structures, indicating that there is increase in antiparallel character. Given that both exhibit the 1630 cm^−1^ peak, and only the antiparallel peak increases in one, we conclude that both the polymorphs contain some amount of parallel structure. The same trend is observed for the 2h fibrils. However, this difference is found to exist in a single fibril, indicating different regions of the same fibril can have different secondary structural organization. Interestingly, we also observe a shift in the beta sheet peak to 1642 cm^−1^ in the 2h fibrils. The absorption frequency of beta sheets is related to their degree of structural order^14^, and a shift to higher wavenumbers thus suggests a more disordered structure compared to oligomers. The later stage fibrils at 6h and 24h exhibit the beta sheet peaks at 1636 cm^−1^ and a more prominent antiparallel peak at 1690 cm^−1^. We do not find any significant heterogeneity in spectra, suggesting that upon maturation, fibrils adopt an ordered, primarily antiparallel structure, consistent with findings from NMR. Given that we do not observe the 1642 cm^−1^ peak in any of the earlier/later aggregates besides 2h fibrils, it is likely that it corresponds to a transient secondary structure. Thus, the 2hr fibrils exhibit a unique secondary structure that is not representative of mature fibrils. That the inherent structural heterogeneity seen in 2h fibrils is not evident in mature fibrils further supports the view that the 2h fibrils represent a transient intermediate that are undergoing structural reorganization. This also underscores that fibrillar morphology does not necessarily represent order, and fibrils can not only have structural disorder, but can also evolve into more ordered structures. Since we do not observe oligomeric species at later stages of aggregation, we conclude that structural reorganization of fibrils can occur without morphological disruption and/or disintegration into smaller aggregates. Taken together, the above results point an aggregation mechanism where early aggregates adopt a distribution of parallel and antiparallel structures, which eventually transforms to antiparallel fibrils. To the best of our knowledge, this parallel to antiparallel transition has never been experimentally identified before. MD simulations have predicted competition between parallel and antiparallel conformations during aggregation of Aβ 16-22^11^; our measurements provide the first experimental evidence of the same. These findings also underscore the significance of spatially resolved spectroscopic measurements: without ability to acquire aggregate specific spectra, it is not possible to determine presence of structural heterogeneity in the structural ensemble, and consequently identify the structural transitions therein. We note that the 1642 cm^−1^ peak can also be attributed to helical conformations. Thus, it is possible that the maturation and reorganization of fibrils proceeds through unconventional secondary structural intermediates. The spectra of 6h and 24h fibrils point towards this possibility: in both we observe a weak band at 1650 cm^−1^, indicating a small amount of residual helical conformations. The presence of helical intermediates before formation of anti-parallel beta structure has been postulated in molecular dynamics (MD) simulations^15^, and our results lend credence to these possibilities.

To test the generality of these structural intermediates, we carried out aggregation of Aβ 16-22 in phosphate buffer, pH 7.4 at 37°C, which better mimics physiological conditions. Within 15 minutes of incubation, oligomers were formed along with small protofibrillar aggregates (Fig 3A). The spectra of these aggregates, shown in Fig. 3B, interestingly do not exhibit the heterogeneity observed for acidic conditions and contain a broad band centered at ~1660 cm^−1^ (Fig. 3B and Fig. S5); second derivative analysis of the mean spectrum shows the presence of underlying bands at 1640 cm^−1^, 1666 cm^−1^ and 1692 cm^−1^ (FIg. 3C), similar to 2h fibrils formed under acidic conditions. This suggests the aggregates contain the same secondary structural components independent of pH, although their relative populations and kinetics can be affected by the aggregation conditions. After 2 hr of incubation, we observe both isolated fibrils and fibril clusters (Fig. 3D and G). Isolated fibrils exhibit a sharp peak at 1624 cm^−1^ (Fig. 3E) with a very small peak at 1690 cm^−1^ (Fig. 3E), indicating predominantly parallel beta structure, whereas fibril clusters show a much more intense 1690 cm^−1^ peak (Fig. 3H and Fig. S5). The relative intensity of these two peaks also varies between different spatial locations, indicating that different regions of the network are at different stages of structural evolution into antiparallel beta structure. This also indicates that the formation of ordered antiparallel structure may be related to integration of isolated fibrils into networks, which is a unique result. After further incubation, at 6h multiple layers of fibrils are observed (Fig. 3J). IR spectra now exhibit more homogeneity and different spectral/structural subclasses are not observed (Fig 3K). The spectra are similar to the 6h fibril spectra under acidic conditions. 24 h fibrils also form a network (Fig. 3M) and the spectra remain the same with two peaks at 1626 cm^−1^ and 1690 cm^−1^ (Fig. 3N), indicating transformation into ordered antiparallel fibrils

**Figure 3:**
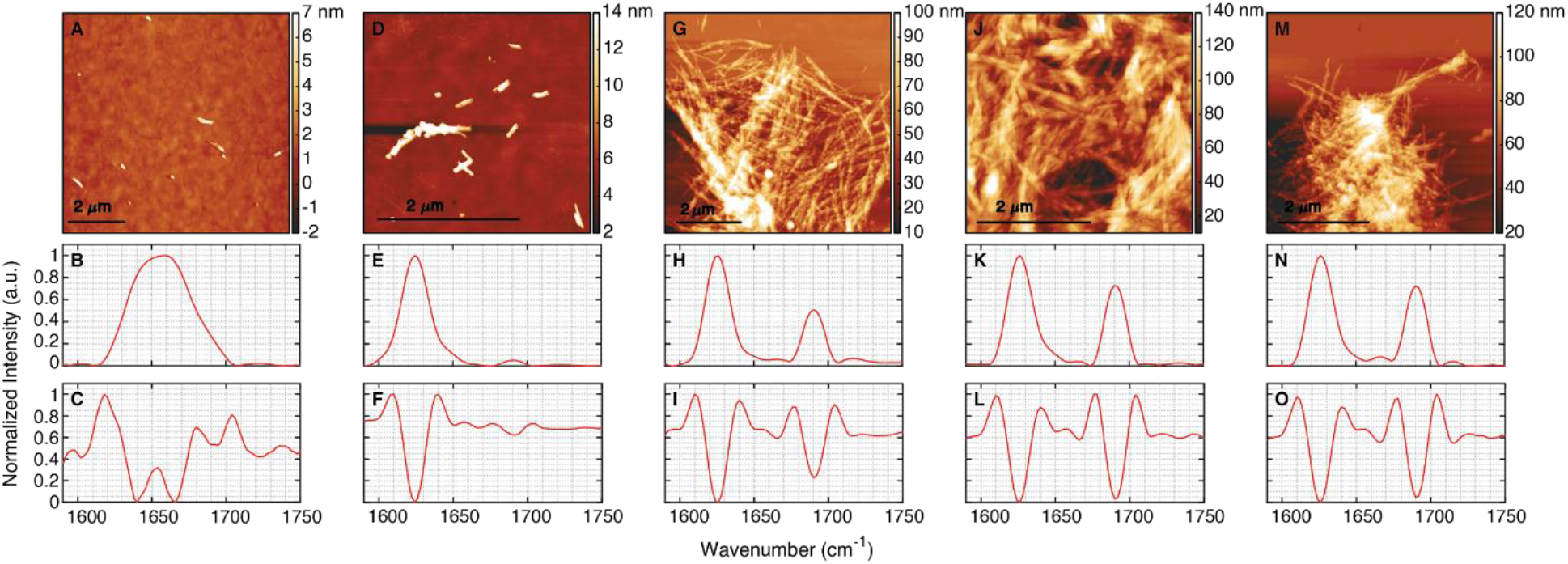
AFM-IR structural study of Aβ 16-22 aggregates in 10 mM phosphate buffer. Top row: AFM images of (A) oligomers produced at 15 min of incubation, (D) isolated fibril and (G) fibril cluster formed after 2 h, (J) fibril cluster after 6 h and (M) 24 h of incubation. Middle row: (B, E, H, K, N) Average IR spectrum of aggregates recorded from corresponding AFM images (top row). Bottom row: (C, F, I, L, O) Second derivatives of corresponding average spectra shown in the middle row.

Taken together, the above results clearly demonstrate that a parallel to antiparallel structural transformation is a fundamental feature in aggregation of Aβ 16-22. Early-stage prefibrillar aggregates do not conform to a singular, uniform secondary structure; rather they contain varying proportions of different beta structures. This structural feature is retained in early-stage fibrils as well, which show clearly different secondary structure between different regions of the fibril. This fibril structure is transient; eventually transitioning into a homogeneous antiparallel structure. The transition appears to be mediated by structural reorganization without loss of morphology. Furthermore, the spectra suggest presence of helical intermediates along with parallel beta structures. These findings imply that a. fibrils are not necessarily structurally ordered and can undergo structural maturation, and b. this maturation can proceed with morphological preservation. The secondary structural evolution of a specific morphological phase in amyloid aggregation has not been addressed or studied in detail, and in absence of polymorphism, fibrillar aggregates are generally perceived to be of a singular structure. Our results offer a unique insight into the maturation of fibrils from the secondary structure perspective and demonstrate that observation of fibrils in morphological characterisations of amyloid aggregates does not guarantee a specific secondary structure; the fibril can undergo spontaneous modifications into a different structure. Aβ 1-40 has been shown to form similar metastable intermediates with different beta sheet structure from mature fibrils^16^, and the presence of such intermediates in other Aβ isoforms has been speculated, but never identified. We demonstrate that presence of such kinetic intermediates is a more general phenomenon. Understanding these structural transitions is critical for elucidating the overall aggregation pathway of larger isoforms of Aβ, since the amyloidogenic core spanning residues 16-21 has been an important target for modulating the aggregation kinetics and developing aggregation inhibitors. This work underscores the necessity of spatially resolved techniques towards investigating the structural reorganization of amyloid peptides and have the potential for significant impact on assessing structural facets of amyloid aggregation.

## Acknowledgement

This work was supported by the National Institutes of Health (Award R35 GM138162).

