## Supplementary Information for "Nanoscale infrared spectroscopy identifies parallel to antiparallel beta sheet transformation of Aβ fibrils"

### Method

#### Aβ 16-22 aggregation

Aβ 16-22 (rPeptide, USA) was completely dissolved in 1,1,1,3,3,3-hexafluoroisopropanol (HFIP) to remove any preformed aggregates. HFIP was evaporated at room temperature (24 ºC) under vacuum. Then the protein was dissolved in two conditions: a. 10 mM HCl and b. 10 mM phosphate buffer, pH 7.4 to prepare 1 mM protein stock. Aggregation was carried out with these two solutions at 37 ºC without any agitation.

#### Sample preparation for AFM-IR experiment

10 μl of aliquots from the aggregation mixture was half diluted and deposited onto ultraflat gold substrates (Platypus Technologies, USA) at 15 min, 2 h, 6 h and 24 h time points. The droplet was incubated on the substrate for 2 min and then rinsed with 500 μl of milli-q water. Then the sample was dried with a gentle stream of nitrogen and kept under vacuum desiccator for 1 h before imaging.

#### AFM-IR experiment

AFM-IR experiments were carried out at room temperature using Anasys Nano IR3 instrument equipped with a mid-IR quantum cascade laser (MIRcat, Daylight solutions). Relative humidity was kept low (less than 5%) by continuous purging of dry air into the instrument during data collection. AFM image and IR data were obtained in tapping mode. The cantilever resonant frequency was 75±15 kHz and the spring constant was 1-7 N/m. Scan speed was kept within a range of 0.5 to 1.0 Hz. During the experiment, a high-resolution AFM image was first recorded, then IR spectra were obtained by moving the tip to the desired locations. Spectral resolution was 2 $cm^{-1}$ and 128 coadditions for each spectral point with 16 coaverages for each spectrum were applied.

#### Data analysis

AFM Images were processed by Gwyddion software. Spectral analysis was carried out in MATLAB software. All spectra were smoothed by applying a (3, 7) Savitzky-Golay filter.

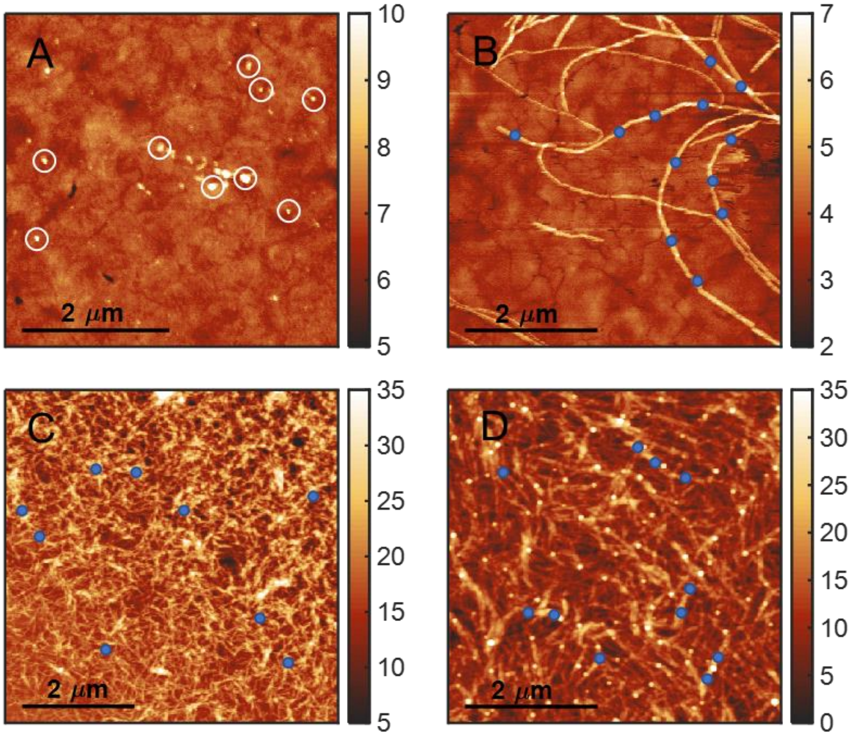


Figure S1. Representative points on AFM topographs showing the locations from where the IR spectra are recorded on Aβ 16-22 aggregates produced in 10 mM HCl. (A) White circles indicate the oligomers where AFM tip was placed for recording IR spectra. (B-D) The blue dots indicate the tip locations for fibril samples.

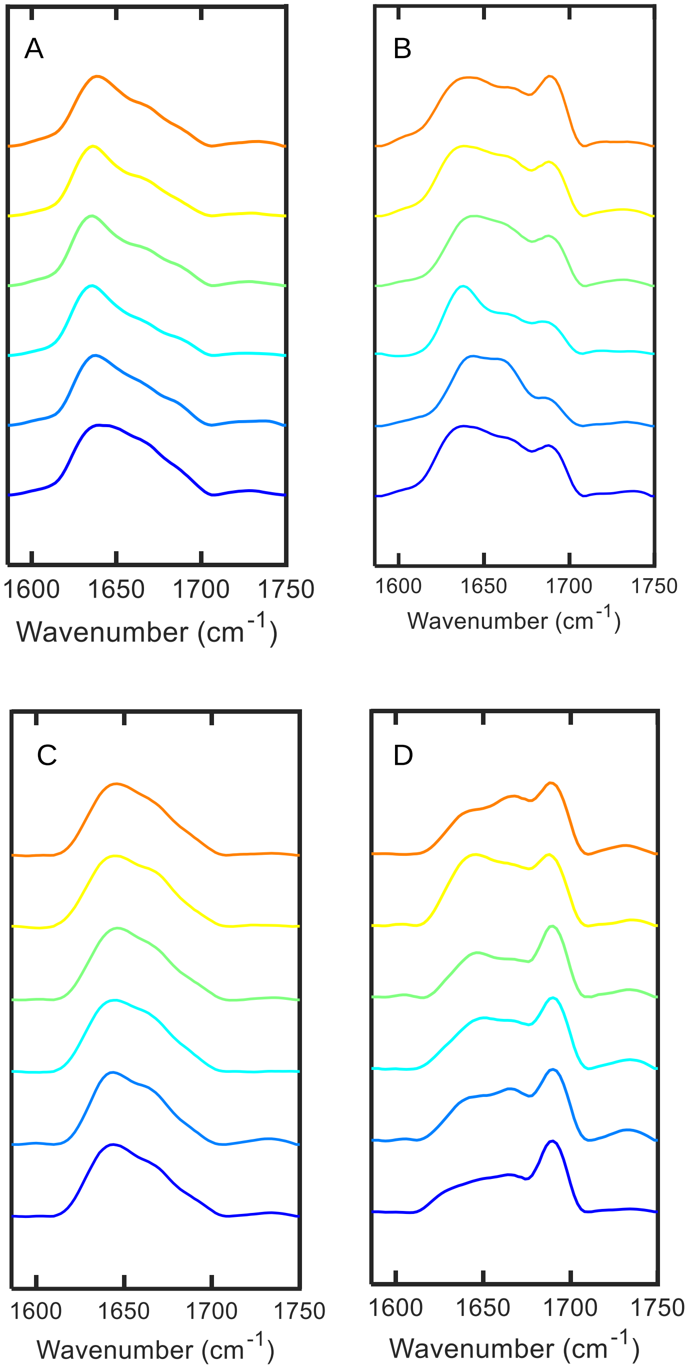


Figure S2. Representative IR spectra obtained from Aβ 16-22 (A-B) oligomers formed after 15 min of incubation in 10 mM HCl and (C-D) fibrils generated after 2 h of incubation in same condition. Two types of amide I bands are observed in both oligomers and fibrils.

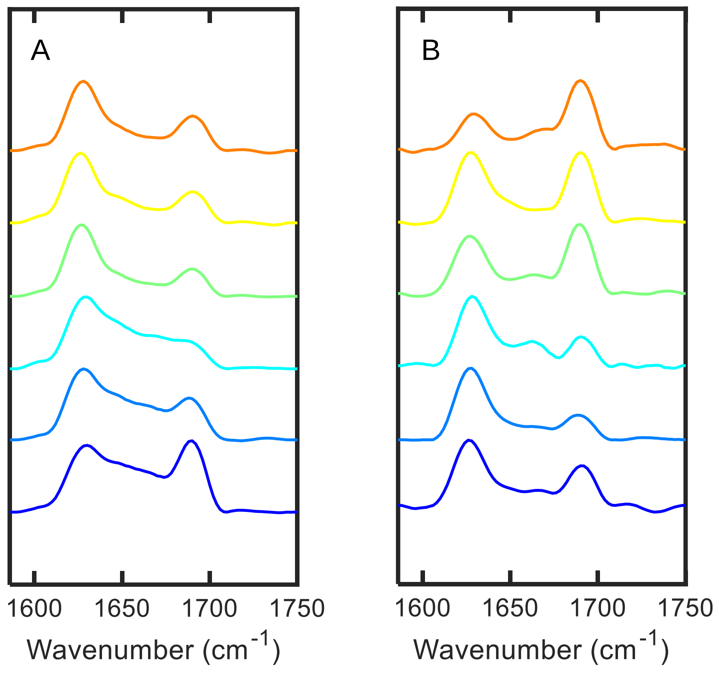


Figure S3. Representative IR spectra recorded from Aβ 16-22 fibrils after (A) 6 h and (B) 24 h incubation in 10 mM HCl.

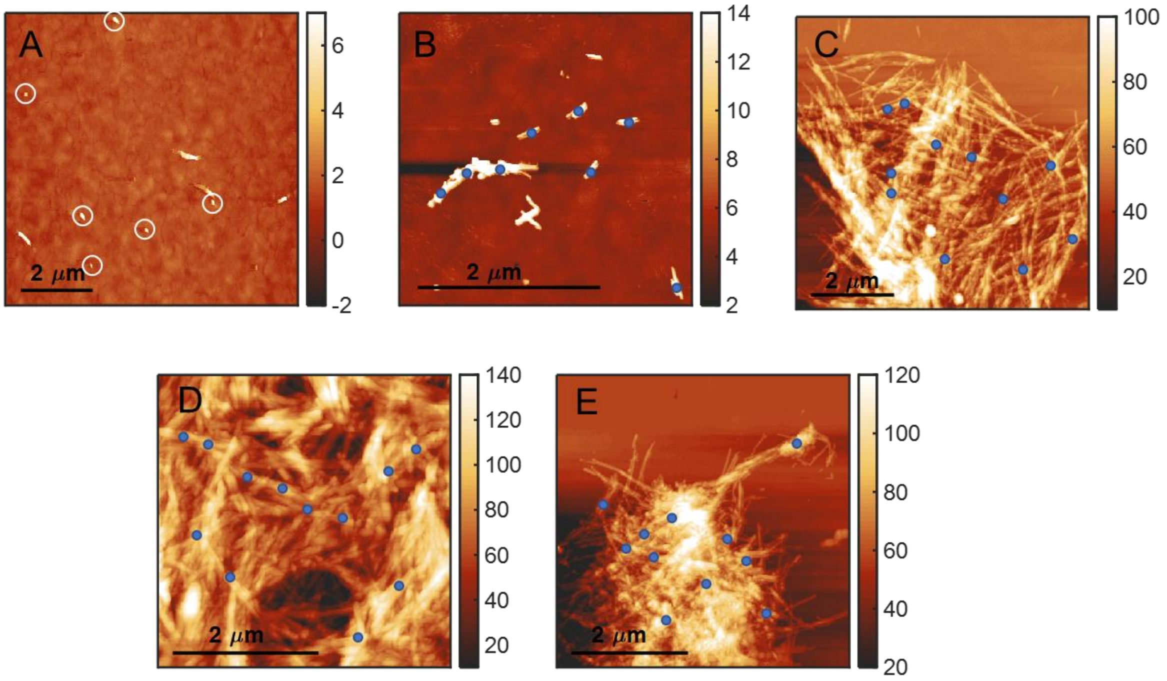


Figure S4. Representative points on AFM topographs showing the locations from where the IR spectra are recorded on Aβ 16-22 aggregates produced in 10 mM phosphate buffer, pH 7.4. (A) White circles indicate the oligomers where AFM tip was placed for recording IR spectra. (B) The blue dots indicate the tip locations for isolated fibril samples at 2 h incubation, (C) fibrils cluster at 2 h, (D) 6 h and (E) 24 h.

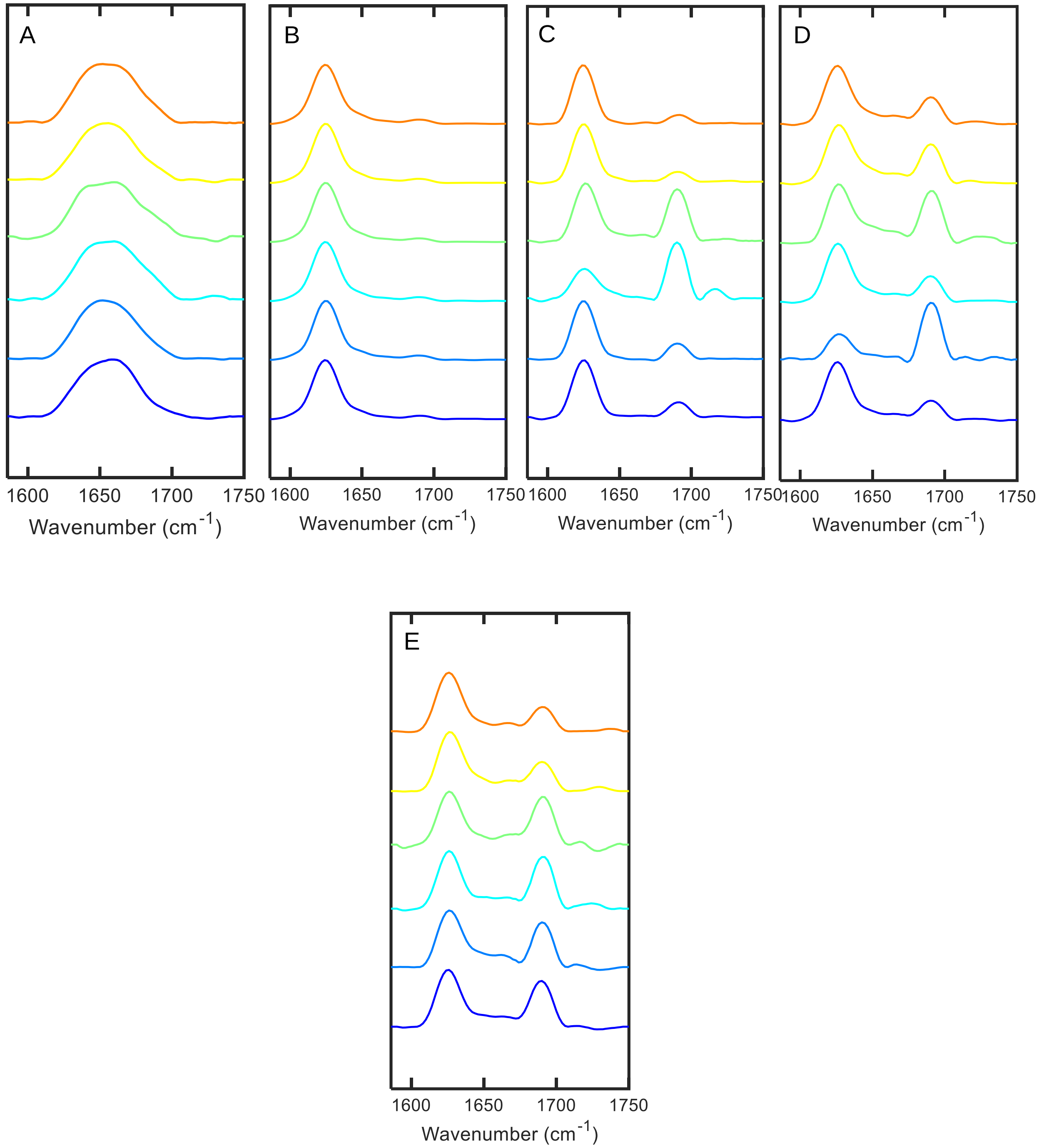


Figure S5: Representative IR spectra recorded from Aβ 16-22 aggregates generated in 10 mM phosphate buffer, pH 7.4 at (A) 15 min, (B) 2 h isolated fibrils, (C) 2 h fibril cluster, (D) 6 h fibrils and (E) 24 h fibrils.
